# Dengue emergence in the temperate Argentinian province of Santa Fe, 2009-2020

**DOI:** 10.1101/2020.08.11.246272

**Authors:** María S. López, Daniela I. Jordan, Evelyn Blatter, Elisabet Walker, Andrea A. Gómez, Gabriela V. Müller, Diego Mendicino, Michael A. Robert, Elizabet L. Estallo

## Abstract

Dengue virus (DENV) transmission occurs primarily in tropical and subtropical climates, but within the last decade it has extended to temperate regions. Santa Fe, a temperate province in Argentina, has experienced an increase in dengue cases and virus circulation since 2009, with the recent 2020 outbreak being the largest in the province to date. The aim of this work is to describe spatio-temporal fluctuations of dengue cases from 2009 to 2020 in Santa Fe Province. The data presented in this work provide a detailed description of DENV transmission for Santa Fe Province by department. These data are useful to assist in investigating drivers of dengue emergence in Santa Fe Province and for developing a better understanding of the drivers and the impacts of ongoing dengue emergence in temperate regions across the world. This work provides data useful for future studies including those investigating socio-ecological, climate, and environmental factors associated with DENV transmission, as well as those investigating other variables related to the biology and the ecology of vector-borne diseases.

## Background & Summary

Dengue virus (DENV serotypes 1-4), which is responsible for dengue fever, is considered one of the most important emerging and reemerging pathogens. Many DENV infections result in mild illness, or even acute flu-like illness, but DENV infections sometimes result in potentially lethal complications such as severe dengue or dengue shock syndrome.

*Aedes aegypti* mosquitoes are the main vectors for DENV as well as for yellow fever, Zika and chikungunya viruses. While DENV transmission is found mainly in tropical and subtropical climates worldwide, the distribution of dengue virus has been expanding into temperate regions in the last decade due to rapid unplanned urbanization, changes in land use, increases in human movement, and changes in climate^1,2,3,4,5^. All of these factors can result in changes in vector distribution and abundance as well as more favorable conditions for DENV transmission^6^. In the last 10 years, DENV’s rapid expansion in temperate regions has generated numerous epidemic events with increasingly larger outbreaks and high incidence rates ^1^.

DENV was eradicated from Argentina in the middle of the past century due in part to successful *Ae. aegypti* control programs^7^; however, during 1997 the first autochthonous transmission in the modern era was reported, and a subsequent outbreak occurred in subtropical northern Argentina^7^. After the reemergence, successive outbreaks appeared in the warmest months and were always closely related to outbreaks in neighboring countries^7^.

During August 2019, the World Health Organization (WHO) warned about the incoming dengue epidemic in the Americas. During the 2019-2020 dengue season, reported cases in the Americas were at their highest ever, with 2,733,635 dengue cases reported through EW 42 of 2019^8^. Neighboring countries of Argentina such as Brazil and Paraguay reported the highest incidences in the region^9,10,11^. As of December 2020, most of Argentina’s provinces (17 of 23) have reported autochthonous cases of dengue since its reemergence^12^.

Santa Fe, a temperate province in central-northeastern Argentina, experienced significant increases in DENV transmission across the last decade with four dengue outbreaks (Figure 1). Santa Fe province is one of the most populated and productive areas of the country. In fact, it features international road connections through the bi-oceanic corridor and the Parana-Paraguay waterway, which gives Santa Fe a privileged geo-strategic location. This central bi-oceanic corridor connects Chile and the Pacific Ocean with Uruguay and the Atlantic Ocean. Santa Fe also connects the southern provinces of Argentina with those of the center and northeast (Figure 1b). Indeed, Santa Fe is a place of passage for land cargo and passengers with Bolivia, Paraguay and Brazil which are neighboring countries with endemic DENV circulation^13^.

**Figure 1.**
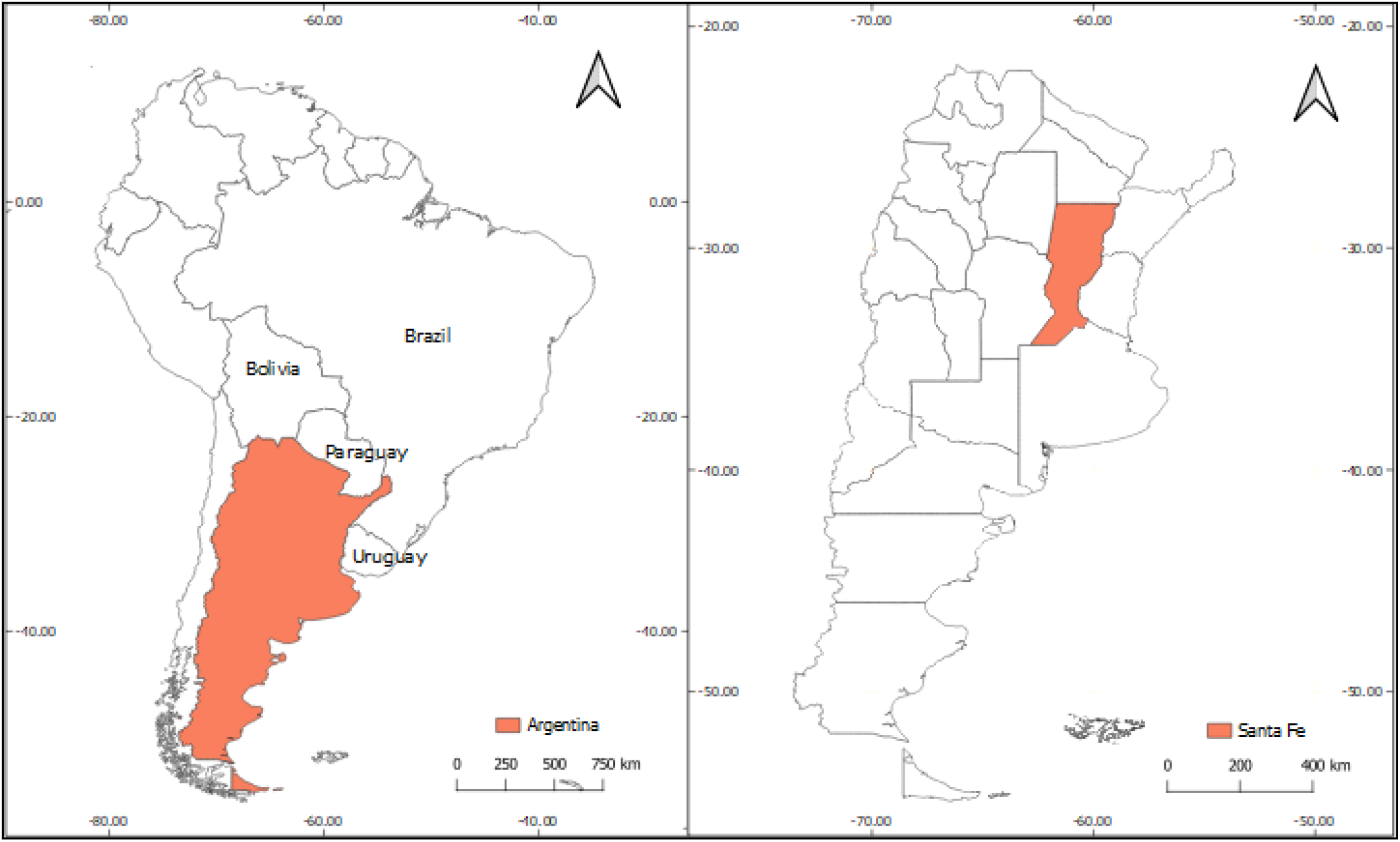
(a) Location of Argentina in South America in relationship to neighboring countries. (b) Map of Argentinian provinces with Santa Fe Province highlighted.

According to the Argentina Ministry of Health (MoH), Santa Fe Province is within the central epidemiological region of the country together with Buenos Aires City and Buenos Aires, Córdoba, and Entre Ríos Provinces, and therefore we will refer to Santa Fe as being within the central region despite Santa Fe being climatically and geographically in central northeastern Argentina. In 2009, Argentina experienced dengue outbreaks in its central region for first time. Since then, dengue cases have been reported each year with the largest number to date occurring in 2020 where more than 50% of the dengue cases of the whole country have been reported in this region. Moreover, in 2020, Santa Fe Province faced its largest dengue epidemic since dengue’s re-introduction in the country despite control efforts of the Health Ministry of Santa Fe (MoS) and the MoH^14^.

The aim of the present work is to perform a detailed description of spatio-temporal fluctuations of dengue cases since 2009 in the temperate Santa Fe Province of Argentina. We also present dengue case distribution and incidence by departments within the province. The database included in this paper is important for future DENV cases studies in the central temperate region of the country, and it is an important source of information for researchers investigating dengue emergence in temperate regions worldwide. Studies of drivers of dengue transmission such as climate change, environmental and meteorological changes, and socio-economic variables will benefit from the data presented in this work as our databases can be combined with other available data to study these and other factors associated with dengue emergence in this region. Studies with mathematical models could utilize this data to investigate previous outbreaks and predict future outbreaks and dengue case occurrence, and this study could be useful for stakeholders in making decisions related to dengue prevention, control, and management at local, national, or even international levels.

## Methods

Dengue epidemics were documented between January 2009 and May 2020 in Santa Fe Province (Figure 1). The region is characterized by a homogeneous geomorphological conformation where the Chaco-Pampeana plain predominates. It consists of a mosaic of wet savannahs and grasslands, subtropical dry forests, gallery forests, shrublands, river inundation plains, and a wide variety of wetlands (e.g., streams, marshes, swamps). The climate in the region is temperate with hot summers and no dry season, according to Köppen-Geiger’s climate classification^15^. The study area presents a latitudinal gradient with maximum temperature and precipitation observed in the north and minimums in the south. The average maximum temperatures in summer range between 32 and 30 °C, and minimum temperatures in winter range between 9 and 3 °C (http://www.smn.gov.ar/serviciosclimaticos). In summer, the monthly precipitation varies between 168 and 136 mm, and in winter it varies between 26 and 13 mm (http://www.smn.gov.ar/serviciosclimaticos). There is more precipitation in the northeast and less in the southwest. The Paraná River is the main waterway and defines the eastern border of the province, along with a complex system of islands, main channels, lagoons, and wetlands. The dynamics of the Paraná River floodplain are strongly shaped by cycles of rises and falls in water levels^16^. The Salado River is another important waterway that crosses the center of the province from west to east flowing into the Paraná River. Santa Fe Province is divided into 19 departments and contains a total population of 3.2 million inhabitants^17^.

Surveillance of arboviruses in Argentina is carried out in an integrated manner, within the surveillance framework for nonspecific acute febrile syndrome and cases that meet specific definitions for each arbovirus. Notifications of suspicious cases are made through the National Health Surveillance System. The MoH, through the National Directorate of Epidemiology and Analysis of the Health Situation, publishes with varying frequency the Integrated Surveillance Bulletin in which the number of total dengue cases are detailed by province (https://www.argentina.gob.ar/salud/epidemiologia/boletinesepidemiologicos).

The data presented here were collected primarily from public health reports provided regularly by the Argentinian National MoH. We identified key data in the reports for characterizing DENV emergence, including the year-to-month cumulative number of dengue probable cases (i.e., with at least one positive laboratory diagnosis) as well as confirmed cases (with two positive laboratory tests), autochthonous (i.e., locally transmitted) and imported (i.e., illness in someone with travel history to a region with arbovirus activity) cases of dengue virus in Argentina. In some years, the reports included information on serotypes of dengue cases and/or regions associated with imported cases (i.e., regions where imported travelers were thought to have acquired the infection) between January 2009 and May 2020. This study does not include suspected and unconfirmed cases (suspected cases have infection symptoms without any laboratory diagnosis).

We analyzed the spatio-temporal fluctuation of DENV cases and identified the areas and periods with the highest incidence of DENV transmission. The cases were reported monthly at a department scale (county scale) between 2009 and 2019. Additionally, cases were reported monthly at the city scale only between January and May 2020. We created a time series of monthly incidence (number of cases per 10,000 inhabitants) to determine the progression of outbreaks during the whole study period. Additionally, we created a time series with the number of cases per epidemiological week (EW) between January and May 2020 to describe the most recent and most important outbreak in the province to date. We also created a dengue incidence map (number of cases per 10,000 inhabitants per department) to facilitate a better understanding of the most affected departments across all years of dengue activity in the province.

### Data Records

The database is publicly available online (via figshare^18^) as a collection of three databases of Santa Fe Province (Argentina) dengue cases. The first file presents data aggregated by province department, for 2009-2020, and contains the total number of cases as well as dengue incidence per 10,000 people. The second file contains the number of cases per month at the province level, divided by imported and autochthonous cases, and the monthly incidence per 10,000 people. The third file presents the Santa Fe cities with the highest number of DENV cases and the corresponding number of cases for each month January-May for the 2020 outbreak. The column headings of the files are as follows.

> YEAR: The year of the date of the entry.
>
> MONTH: The month of the date of the entry.
>
> AUTO_CASES: Number of autochthonous cases.
>
> IMP_CASES: Number of imported cases.
>
> TOTAL 2009-2020: The sum of the number of cases that occurred between 2009 and 2020.POP_SIZE: Population size based on estimates of the National Institute of Statistics and Census (INDEC, https://www.indec.gob.ar/indec/web/Nivel4-Tema-2-24-119).
>
> POP_SIZE_2009: Population size in 2009 from INDEC.
>
> INCIDENCE 2009-2020: The total incidence (per 10,000 people) of dengue between 2009-2020 calculated using the 2009 population size from INDEC.AUTO_INC: Incidence of autochthonous cases (per 10,000 people).
>
> IMP_INC: Incidence of imported cases (per 10,000 people).
>
> EPID_WEEK: The epidemiological week described by MoH bulletin of the given date.
>
> AUTO_IMP_CASES: The total number of autochthonous and imported cases. DEPARTMENT: Santa Fe Province department name.
>
> CITY: Name of the city.
>
> INCIDENCE: Total incidence by Santa Fe Province department (per 10,000 people).

A total of 6,454 DENV cases were reported in Santa Fe province between January 2009 and May 2020 (6,209 autochthonous, 245 imported). Figure 2 and Table 1 characterize dengue emergence in the studied period. Four dengue outbreaks have been reported in Santa Fe Province since 2009, with the most severe one reported during 2020 with 4521 dengue cases, 4457 of which were autochthonous and 64 of which were imported (total incidence of 12.78 per 10,000 people). As shown in Figure 2 during 2016 and 2019 the province also experienced outbreaks, with 2016 being the more intense one of these two. During the 2016 outbreak 1014 dengue cases were reported with 929 autochthonous and 85 imported cases (total incidence of 2.96 per 10,000 people). During 2019, 484 dengue cases were reported with 467 autochthonous and 17 imported cases (total incidence of 1.37 per 10,000 people). During the first outbreak that reached central Argentina in 2009, 154 dengue cases were reported, where 120 were autochthonous and 34 were imported cases (total incidence of 0.47 per 10,000 people) (Table 1). Imported dengue cases originated mainly from tropical countries where dengue fever is endemic, as well as from the northern region of Argentina, although there were also imported cases originating in temperate countries such as Uruguay (Table 2). Although DENV circulation typically occurs each season between January and May, ten autochthonous dengue cases have been reported in Santa Fe between June and December: during August 2009 (N = 1), July 2011 (N = 1), October (N = 1) and December 2013 (N = 1), November 2016 (N = 2), and June 2018 (N = 4).

**Figure 2.**
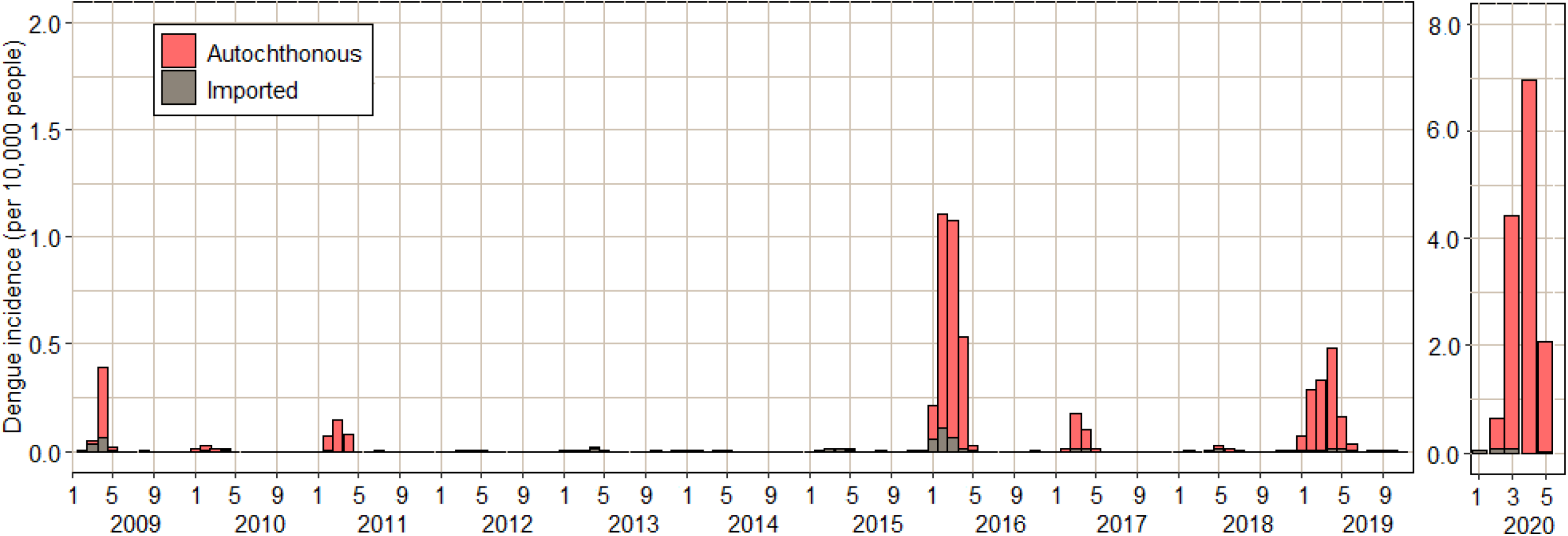
Incidence of imported and autochthonous dengue cases each month between January 2009–May 2020. Incidence is calculated as the number of cases per 10,000 inhabitants of Santa Fe Province. Population estimates were obtained for each year from projections of the National Institute of Statistics and Censuses.

Figure 3 shows the incidence of dengue by departments (i.e., counties) of the Province of Santa Fe during January 2009 to May 2020. Dengue incidence in Santa Fe departments was clearly highest in the northeast area of the province that borders with Chaco and Corrientes Provinces located northeast of the country. The predominant serotype that circulated among all outbreaks was DENV-1 although all four DENV serotypes were detected between 2009-2020 (Table 1). During the 2020 outbreak, 56.3% of the cases were detected as DENV-1 (488 cases), 42.98% as DENV-4 (371 cases) and 0.46% as DENV-2 (4 cases) (Table 1). DENV-4 was widely distributed in the province, DENV-2 was reported only in Rosario department and DENV-3 was not reported during this outbreak. Between EW 9 and EW 17, five deaths were reported (0.11 % of total cases in the province). Table 2 shows the most affected cities during the 2020 outbreak, where almost 80% of cases were reported.

**Figure 3.**
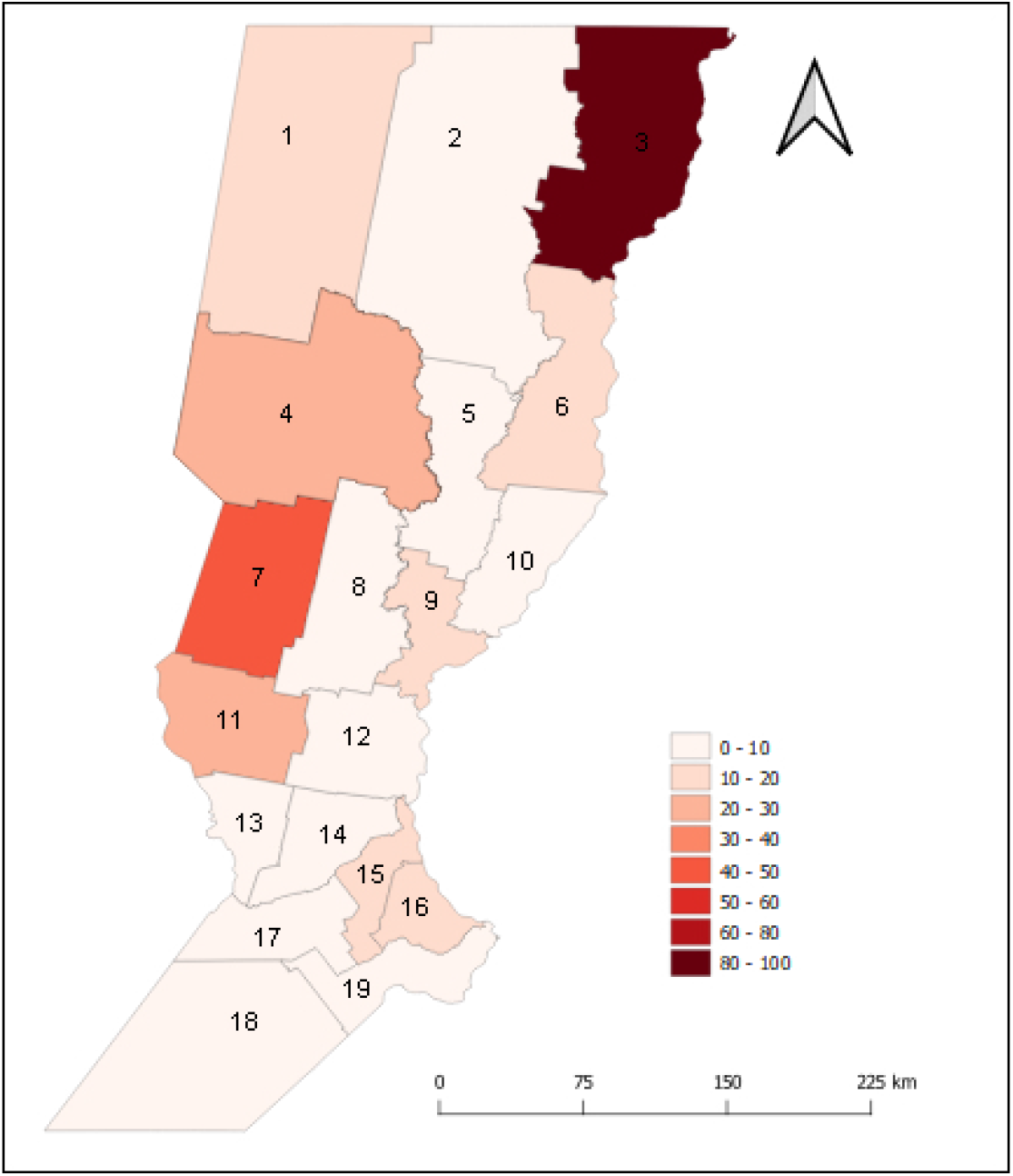
Total incidence of imported and autochthonous dengue cases between January 2009–May 2020 by department. Incidence is calculated as the number of cases per 10,000 inhabitants by department of Santa Fe Province. 1. 9 de Julio, 2 Vera, 3 General Obligado, 4 San Cristóbal, 5 San Justo, 6 San Javier, 7 Castellanos, 8 Las Colonias, 9 La Capital, 10 Garay, 11 San Martin, 12 San Jerónimo, 13 Belgrano, 14 Iriondo, 15 San Lorenzo, 16 Rosario, 17 Caseros, 18 General López, 19 Constitución.

### Technical Validation

The National Health Surveillance System operated by the MoH reports suspected dengue infections at either private or public clinical sites. The Central Reference Laboratory of the City of Santa Fe and the Ambulatory Medical Center of Rosario City (both in Santa Fe Province) analyzed samples. The selection of the diagnostic methods by either indirect or direct methods is made based on the number of days of the evolution of the symptoms (https://www.argentina.gob.ar/salud/epidemiologia/boletinesepidemiologicos). The Santa Fe laboratories sent a subset of samples to the National Reference Laboratory (INEVH-Maiztegui) in the City of Pergamino, Buenos Aires Province, where the diagnostic methods are carried out by viral isolation and neutralization with a flavivirus panel.

The epidemiological information is published online in Spanish at the scale of provinces by the MoH through epidemiological bulletins (https://www.argentina.gob.ar/salud/epidemiologia/boletinesepidemiologicos). For the purpose of this study, we requested additional case data at the department level from the Santa Fe Province ministry of health. We also requested data at the city level for January-May 2020 from the MoH. This data was made available in part due to the public information open access national law (N° 27275), with the protection of personal data regulated by the National Law N° 25326.

### Usage Notes

The data presented herein show the increases of DENV cases and DENV transmission in the last decade in Santa Fe Province, highlighting the 2020 outbreak as it is most important outbreak to date. This data also emphasizes the importance of this area in the ongoing dengue emergence in the Southern Cone because of its geographic location relative to other provinces and neighboring countries were DENV has endemic circulation.

Argentina experienced its most important season of dengue to date in 2020, when 69.5% of the national territory (16 of 23 provinces) were affected by the dengue outbreak. The dengue outbreak in Santa Fe Province during 2020 was four times larger than the 2016 outbreak. The department of General Obligado, in the northeast region of the province, presented co-circulation of DENV 1 and 4. This area also connects to northern provinces of the country that border the neighboring countries of Brazil and Paraguay were dengue regularly has high incidence, suggesting southern spread of dengue transmission is being driven by importation from neighboring countries. In addition to increases in overall transmission, concern over dengue transmission in the region is mounting due to the presence of multiple serotypes co-circulating, which presents a risk for increased severity of dengue due to reactions among serotypes (https://www.paho.org/en/topics/dengue). In 2020, areas of southern Brazil reported co-circulation of DENV 1, 2, and 4 until EW 23^10^. In Paraguay, cases of DENV 1, 2, and 4 were identified^11^. The co-circulation of a similar number of confirmed cases of DENV 1 and 4 (Table 1) in Santa Fe Province increases the potential for severe dengue cases(https://www.paho.org/en/topics/dengue). Co-circulation of more than one serotype and the increases in dengue severity caused by this is a high concern for Santa Fe Province, and other temperate areas where dengue is actively emerging.

Global predictions for dengue distribution emphasize that transmission is most likely to occur in tropical regions, with areas of greatest risk in Asia and The Americas^19^. Risk assessments such as these often consider factors such as climate, human movement, and distributions of vectors. To assess risk in Santa Fe province, the data presented here could be coupled with climate data to better understand the relationships between changes in climate and meteorological anomalies and ongoing dengue emergence^19^. Another key factor in dengue transmission is human movement^3,19^, and the data herein could be utilized to investigate how human movement patterns in Santa Fe Province and in Argentina could potentially be driving recent dengue activity. Furthermore, a global *Ae. aegypti* and *Ae. albopinctus* distribution study shows that among the most important predictors of *Aedes* populations is vegetation indices^20^. Socio-ecological behavior associated with cities and rapid urbanization generate appropriate places for mosquito development^6^, and unplanned growth can lead to the lack of essential services such as garbage depots, insufficient health systems, and comprehensive entomological surveillance^7^. Coupled with vegetation indices and sociodemographic data, the data presented here could be used to investigate the role that urbanization and changes in socioeconomical dynamics have played in dengue transmission. Finally, the 2020 outbreak is in unique context due to the COVID-19 pandemic, during which people in Argentina have had mandatory quarantines. It is possible that the dengue epidemic in 2020 was particularly severe in part due to the increased time that people are spending in their homes, which potentially increases exposure to *Ae. aegypti* mosquitoes^21^, as well as reductions in mosquito surveillance and control that have occurred during this time^22^. The data presented here will be useful for understanding the role that the COVID-19 pandemic has had on the dynamics of other infectious diseases by providing one case study of a region that has experienced increases in vector-borne disease transmission during the COVID-19 pandemic.

The Argentina dengue control plan is based on integrated strategies for diminishing the vector population. Therefore, it is necessary to focus researchers on spatio-temporal dynamics of DENV transmission to improve entomological surveillance. The data presented in this work provide a detailed description of DENV transmission for Santa Fe Province by department to highlight the recent and ongoing emergence of dengue in the province. This information together with other works in the temperate Argentina^2,21^ will be useful in better understanding the impact of dengue emergence and reemergence in other areas of the world. Indeed, this work can be combined with other existing data sets to contribute to future studies including those aimed at investigating socio-ecological, climate, and environmental factors associated with dengue emergence, as well as those aimed at understanding the influence of other variables related to the biology and the ecology of vector-borne diseases.

## Acknowledgements

The data of dengue cases were granted by Ministry of Health and meteorological data by National Meteorological Service of Argentina (SMN). MSL, EW, AG, GVM and ELE are members of Consejo Nacional de Investigaciones Científicas y Tecnológicas (CONICET) from Argentina.

## Author Contributions

Conceived of the Research: ELE

Extracted and Processed Data: MSL, DJ, EB

Analyzed Data: MSL, DJ

Generated Figures and Tables: MSL, EW Wrote the

Manuscript: ELE, MSL

Revised and Reviewed the Manuscript: MAR, ELE, MSL, DM, GM, AG

## Competing Interests

The authors declare no competing interests.

## Figure and Tables Legends

Table 1. Total incidence of confirmed and probable cases, number of cases confirmed by serotypes, confirmed DENV serotypes, and origin of imported cases. Incidence is calculated as the number of cases per 10,000 inhabitants of Santa Fe Province. The population sizes used for incidence counts are those obtained from the 2010 census.

Table 2.Cities with the highest number of dengue cases in the 2020 outbreak in the Santa Fe Province. Incidence is calculated as the number of cases per 10,000 inhabitants.

## Notes

### Competing Interest Statement

The authors have declared no competing interest.

## References

1. Velasquez-Serra, G. et al. Arbovirosis De Importancia En Las Regiones Tropicales (Centro de Investigación y Desarrollo Profesional CIDEPRO Press, 2020)

2. Robert, M. A. et al. Arbovirus emergence in temperate climates: the case of Córdoba, Argentina, 2009–2018. Sci. Data. 6, (2019).

3. Estallo, E. L. et al. Spatio-temporal dynamics of dengue 2009 outbreak in Córdoba City, Argentina. Acta Trop. 136, 129–136 (2014).

4. Estallo, E. L. et al. Modelling the distribution of the vector Aedes aegypti in a central Argentine city. Med.Vet.Ent. 32, (2018).

5. Confalonieri, U. & Quintão, A. F. Vulnerabilidade a mudança climática na America Latina: intrumentos regionais para a adaptação no sector saúde In Vulnerabilidade à mudança climática na América Latina: instrumentos regionais para a adaptação no setor saúde (Belo Horizonte, 2016).

6. Bernardini Zambrini, D. Neglected lessons from the 2009 dengue epidemic in Argentina. Rev. Saú. Púb. 45, 428–31 (2011).

7. Martino, O. & Weissenbacher, M. Historia natural de enfermedades emergentes y reemergentes en la Argentina: Zika, chikungunya y dengue (2016-2017). Pren. Méd. Argent. 103, 365–375 (2017).

8. Pan American Health Organization. Actualización Epidemiológica Dengue. Noviembre de 2019, https://www.paho.org/es/documentos/actualizacion-epidemiologica-dengue-11-noviembre-2019. (2019).

9. Ministerio de Salud de Argentina. Boletín Integrado de Vigilancia. Número 484. Semana epidemiológica 5 https://www.argentina.gob.ar/sites/default/files/biv_484_edicion_semanal.pdf (2020).

10. Ministerio de Salud de Brazil. Secretaria de Vigilancia en Salud. Boletin Epidemiológico 24. https://antigo.saude.gov.br/images/pdf/2020/June/16/Boletim-epidemiologico-SVS-24-final.pdf (2020).

11. Ministerio de Salud Pública y Bienestar Social de la República del Paraguay. http://vigisalud.gov.py/page/#vista_boletines_dpto.html (2020).

12. Ministerio de Salud de Argentina. Boletín Integrado de Vigilancia. Número 499. Semana epidemiológica 23 https://www.argentina.gob.ar/sites/default/files/biv_499_se23.pdf (2020).

13. Province of Santa Fe, Argentine Republic. Strategies and Capacities for a Competitive Global Insertion. Ministry of Government and Reform of the State of the Province of Santa Fe, https://www.santafe.gov.ar/index.php/web/content/download/243244/1281334/file/Santa%20Fe%20-%20Strategies%20and%20Capacities%20for%20a%20Competitive%20Global%20Insertion.pdf (2016).

14. Ministerio de Salud de Argentina. Plan Nacional de Prevención y Control del Dengue y la Fiebre Amarilla, https://www.fundacionfemeba.org.ar/blog/farmacologia-7/post/plan-nacional-de-prevencion-y-control-del-dengue-y-la-fiebre-amarilla-42755 (2009).

15. Peel, M., Finlayson, B. L. & Mcmahon, T. A. Updated world map of the Köppen-Geiger climate classification. Hydrol Earth Syst Sci. 4, 1633–1644 (2007).

16. Burkart, R., Bárbaro N. O., Sánchez R. O. & Gómez, D. A. Ecorregiones De La Argentina. (APN, PRODIA, 1999).

17. Provincia de Santa Fe. Población según Censo Nacional de Población 2010. https://www.santafe.gov.ar/index.php/web/Estructura-de-Gobierno/Ministerios/Economia/Secretaria-de-Planificacion-y-Politica-Economica/Direccion-Provincial-del-Instituto-Provincial-de-Estadistica-y-Censos-de-la-Provincia-de-Santa-Fe/ESTADISTICAS/Censos/Poblacion/Censo-Nacional-de-Poblacion-y-Vivienda-2010/Estadisticas-por-Dpto.-y-Pcia/Poblacion/Poblacion-segun-Censo-Nacional-de-Poblacion-2010.-Provincia-Santa-Fe (2010).

18. López, M. S. et al. Dengue arbovirus affecting temperate Argentina province for more than a decade (2009-2020). figshare. https://figshare.com/s/6d5256dbff440927827b (2020).

19. Bhatt, S. et al. The global distribution and burden of dengue. Nature. 496, 504–507. (2013).

20. Kraemer, M. et al. The global distribution of the arbovirus vectors Aedes aegypti and Ae. albopictus. Ecology, Epidemiology and global health. eLife 4:e08347 (2015).

21. Robert, M. A., Stewart-Ibarra, A. M & Estallo, E. L. Climate change and viral emergence: evidence from Aedes-borne arboviruses. Curr. Opin. Virol. 40, 41–47 (2020).

22. Nacher M. et al. Simultaneous dengue and COVID-19 epidemics: Difficult days ahead? PLoS Negl Trop Dis 14(8): e0008426 (2020)

