## Supplementary material for "Dengue emergence in the temperate Argentinian province of Santa Fe, 2009-2020": Table 1

| YEAR | Total incidence of confirmed and probable cases | N° cases identified by serotypes | DENV Serotypes Detected | Origins of imported cases |
|---|---|---|---|---|
| 2009 | 0.472 | 88 | DENV-1 (N = 88) | Argentina provinces: Chaco, Córdoba, Santiago del Estero |
| 2010 | 0.058 | 6 | DENV-1 (N = 4), DENV-2 (N =1), DENV-4 (N = 1) | Brazil |
| 2011 | 0.203 | 32 | DENV-1 (N = 30), DENV-2 (n = 2) | No information |
| 2012 | 0.009 | 2 | DENV-1 (N = 1), DENV-4 (N = 1) | Brazil |
| 2013 | 0.047 | 14 | DENV-1 (N = 3), DENV-2 (N =5), DENV-4 (N = 6) | Brazil, Paraguay |
| 2014 | 0.014 | 3 | DENV-1 (N = 3) | Brazil, Dominican Republic |
| 2015 | 0.05 | 15 | DENV-1 (N = 15) | Brazil, Ecuador. Argentina provinces: Formosa and Misiones |
| 2016 | 2.96 | 277 | DENV-1 (N = 267), DENV-4 (N = 10) | Brazil, Paraguay, Thailand, Uruguay. Argentina provinces: Buenos Aires, Chaco, Córdoba, Corrientes, Entre Ríos , Formosa, Mendoza, Misiones |
| 2017 | 0.304 | 76 | DENV-1 (N = 76) | Brazil, Chile. Argentina provinces: Buenos Aires, Córdoba, Jujuy |
| 2018 | 0.067 | 8 | DENV-1 (N = 7), DENV-3 (N = 1) | Argentina provinces: Formosa |
| 2019 | 1.379 | 75 | DENV-1 (N = 71), DENV-2 (N =3), DENV-4 (N = 1) | Brazil, Indonesia, México. Argentina provinces: Misiones, Salta |
| 2020 | 12.781 | 863 | DENV-1 (N = 488), DENV-2 (N = 4), DENV-4 (N = 371) | Brazil, México, Paraguay. Argentina provinces: Chaco, Córdoba, Corrientes, Formosa, La Rioja, Misiones, Salta. |
