## Supplementary material for "Dengue emergence in the temperate Argentinian province of Santa Fe, 2009-2020": Table 2

| Cities | Total of confirmed and probable cases | Population of the cities | Total incidence (per 10000 people) of confirmed and probable cases | Latitude | Longitude |
|---|---|---|---|---|---|
| Rosario | 1212 | 948.312 | 12.78 | 32° 57’ 25.51’’ S | 60° 41’ 36.84’’ O |
| Reconquista | 1096 | 73.293 | 149.53 | 29° 08’ 39.89’’ S | 59° 38’ 55.18’’ O |
| Rafaela | 629 | 92.945 | 67.67 | 31° 15’ 08.58’’ S | 61° 29’ 29.49’’ O |
| Avellaneda | 280 | 25.995 | 107.71 | 29° 07’ 03.79’’ S | 59° 39’ 29.98’’ O |
| Santa Fe | 239 | 391.231 | 6.10 | 31° 36’ 39.07’’ S | 60° 41’ 49.36’’ O |
| San Jorge | 147 | 18.056 | 81.41 | 31° 53’ 49.23’’ S | 61° 51’ 35.16’’ O |
| Fray Luis Beltran | 102 | 15.389 | 66.28 | 32° 47’ 00.45’’ S | 60° 43’ 58.02’’ O |
